# The challenges of estimating the distribution of flight heights from telemetry or altimetry data

**DOI:** 10.1101/751271

**Authors:** Guillaume Péron, Justin M. Calabrese, Olivier Duriez, Christen H. Fleming, Ruth García-Jiménez, Alison Johnston, Sergio Lambertucci, Kamran Safi, Emily L.C. Shepard

## Abstract

**Background:** Global positioning systems (GPS) and altimeters are increasingly used to monitor vertical space use by aerial species, a key aspect of their niche that we need to know to understand their ecology and conservation needs, and to manage our own use of the airspace. However, there are various sources of error in flight height data (“height” above ground, as opposed to “altitude” above a reference like the sea level): vertical error from the devices themselves, error in the ground elevation below the tracked animals, and error in the horizontal position of the animals and thus the predicted ground elevation below them.

**Methods:** We used controlled field trials, simulations, and the reanalysis of raptor case studies with state-space models to illustrate the effect of improper error management.

**Results:** Errors of a magnitude of 20 meters appear in benign conditions (expected to be larger in more challenging context). These errors distort the shape of the distribution of flight heights, inflate the variance in flight height, bias behavioural state assignments, correlations with environmental covariates, and airspace management recommendations. Improper data filters such as removing all negative recorded flight height records introduce several biases in the remaining dataset, and preclude the opportunity to leverage unambiguous errors to help with model fitting. Analyses that ignore the variance around the mean flight height, e.g., those based on linear models of flight height, and those that ignore the variance inflation caused by telemetry errors, lead to incorrect inferences.

**Conclusion:** The state-space modelling framework, now in widespread use by ecologists and increasingly often automatically implemented within on-board GPS data processing algorithms, makes it possible to fit flight models directly to raw flight height records, with minimal data pre-selection, and to analyse the full distribution of flight heights, not just the mean. In addition to basic research about aerial niches, behaviour quantification, and environmental interactions, we highlight the applied relevance of our recommendations for airspace management and the conservation of aerial wildlife.

## Introduction

Describing the distribution of animals in environmental space is fundamental to understanding their resource requirements, cognitive processes, energetic strategies, and ecological characteristics. The distribution of animals in horizontal space has dominated ecological studies (Nathan et al. 2008), however the vertical dimension is also important for flying animals, and for that matter also diving and tree-climbing animals (Weimerskirch et al. 2005, Kunz et al. 2007, Bishop et al. 2015, Liechti et al. 2018). For example, flight height data could help documenting the vertical niche and community ecology of aerial foragers (Arlettaz et al. 1997, Siemers and Schnitzler 2004). Flight height data quantify the behaviour of flying animals and their flight strategies (Pirotta et al. 2018, Murgatroyd et al. 2018), and their relationships with environmental factors (e.g., Péron et al. 2017). From an applied perspective, we need an accurate, error-free description of the distribution of birds and other animals in the aerosphere to avoid collisions with man-made structures, which is key to aircraft safety and animal conservation, in the current context of increasing human encroachment into the airspace (Lambertucci et al. 2015, Davy et al. 2017).

However, monitoring vertical airspace use by wildlife remains challenging. Ground-based surveys are limited in their field of vision and time window. Airborne monitoring (e.g., from glider planes) is logistically challenging and constrained by weather conditions. Radar-based methodologies are not usually specific enough to assign records to species (but see Zaugg et al. 2008, Dokter et al. 2013). Animal-borne tracking methodologies such as global positioning systems (GPS) and altimeters have therefore become popular to monitor flying species (López-López 2016). They record data even when the animals are out of sight for ground-based observers, over extensive, potentially uninterrupted periods of time, and with no uncertainty about which species or individuals are being monitored. For example, we can record raptors soaring over the high sea at night (Duriez et al. 2018). However, the data that GPS and altimeters record are not error-free (D’Eon et al. 2002, Frair et al. 2004, Jerde and Visscher 2005, Brost et al. 2015). Errors are particularly evident in the vertical axis because there are unpassable barriers, e.g., the ground. Usually, a few unambiguously erroneous positions are recorded beyond these barriers (Katzner et al. 2012, Ross-Smith et al. 2016, Weimerskirch et al. 2016, Péron et al. 2017, Krone and Treu 2018, Roeleke et al. 2018).

Most of the research into ways to deal with sampling errors in positioning data has focused on horizontal animal movement (Freitas et al. 2008, Albertsen et al. 2015, Brost et al. 2015, Fleming et al. 2016). There is very little guidance for ecologists about the challenges specific to vertical space use data (Poessel et al. 2018). Many practitioners consider that vertical movement data need to be “filtered” before analysis, i.e., they discard some records before proceeding with the analysis. They may discard records that are too far from preceding ones (as often done for horizontal data; Freitas et al., 2008), too far beyond impassable barriers (Katzner et al. 2012, Krone and Treu 2018), or obtained from an unreliable configuration of the GPS satellite network (Poessel et al., 2018). Instead of discarding the more erroneous records, researchers have also sometimes chosen to reset them to plausible values (Weimerskirch et al. 2016, Roeleke et al. 2018). However, when applied improperly, such filters can have undesirable consequences. We start by reviewing the sources of error in GPS and altimeter flight height data (Part 1). In Part 2, we reanalyse case studies into the flight height of three raptor species (Péron et al. 2017), and complement them with novel data from controlled field trials and from simulations, in order to illustrate the stakes of proper error-handling in vertical airspace use data.

## Part 1: Review of the sources of error in flight height data from GPS and altimeters

Throughout we refer to flight height *h*, which is the distance to the ground below the bird, different from flight altitude *z*. The flight altitude denotes the distance to a reference altitude, often the ellipsoid, i.e., a geometrically perfect (but simplistic) model of the sea level. Alternatively, some GPS units may provide the altitude relative to the empirical sea level measured at a reference point over a reference period (e.g., in France the “NGF-IGN 1969” norm means that altitude is measured relative to the mean sea level in the port of Marseille between 1884 and 1896), or relative to the geoid, which is a model of the sea level if it was only influenced by the local gravitational field and the rotation of the Earth (Fowler 2005). There are databases and simple formulae to convert from one system of reference to another, but this nevertheless represents a first potential source of error in flight height data.

Flight height above the ground is computed as *h* = *z− z_DEM_* (*x, y*), where *z_DEM_* (*x, y*) is the ground altitude predicted by a digital elevation model (DEM) at the recorded horizontal position (*x, y*), in the same system of reference as *z*. Errors in *h* can then be caused by errors in any of the three components: *z, z_DEM_*, or (*x, y*) (Fig. 1). Importantly, depending on the application, researchers might want to study *z* not *h* (Pirotta et al. 2018, Murgatroyd et al. 2018). In the list below, sources of error #3-#5 do not influence *z*.

**Fig. 1:**
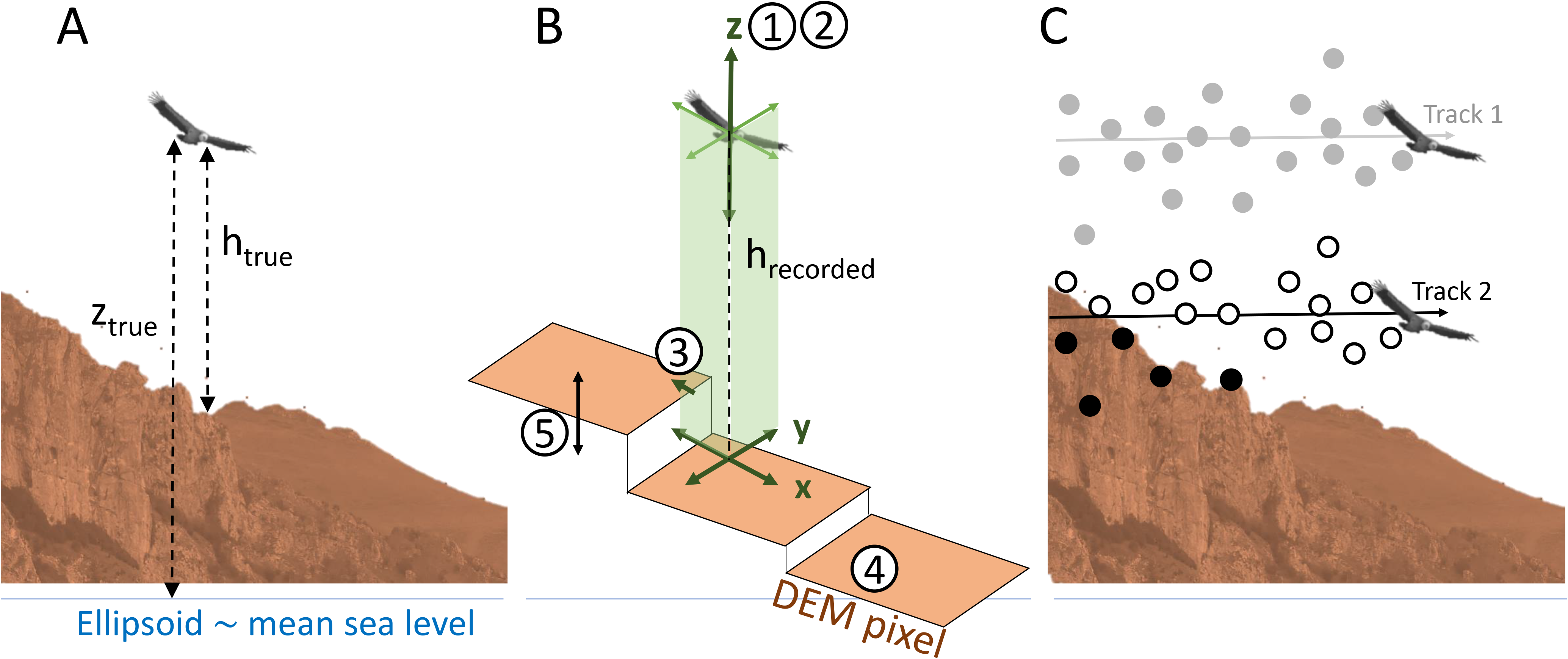
Illustration of the difference between *true* and *recorded* flight height. A: True flight height above ground (*h_true_*), and true elevation above ellipsoid (*z_true_*). B: Adding the five sources of error, with circled numbers referring to headings in Part 1. DEM stands for Digital Elevation Model. C: Two tracks with the same amount of error. The bird of track 1 is flying high so all the recorded flight height data remain positive despite the errors. The bird of track 2 is flying low, so some of the recorded data fall below the digital elevation model.

### 1. Error in *z* when *z* is given by a GPS

If recorded by a GPS, z is affected by the “user equivalent range error” (UERE) and the “vertical dilution of precision” (VDOP) (Parkinson and Spilker 1996, Sanz Subirana et al. 2013).

The UERE stems from diffusion and diffraction in the atmosphere, reflection from obstacles, and receiver noise (Parkinson and Spilker 1996, Sanz Subirana et al. 2013). The acronym UERE usually directly refers to the root mean squared error, but here we will use the notation σ_UERE_ instead. ^σ^UERE ^is^ usually in the order of a few meters and considered constant over time for a given device. Some GPS manufacturers specify the horizontal σ_UERE_, or alternatively it can be estimated from the data (Johnson et al. 2008). The σ_UERE_ is however reputedly larger in the vertical axis than the horizontal axes (D’Eon et al. 2002, Bouten et al. 2013), meaning that manufacturer-provided σ_UERE_ should be considered conservative for vertical applications and should be used with appropriate caution.

The vertical position dilution of precision factor (VDOP) quantifies the effect of changes in the size and spatial configuration of the available satellite network on the precision of GPS records (Parkinson & Spilker, 1996; Sanz Subirana et al., 2013; Fig. A1). The more satellites are available and the more evenly spread apart they are, the more reliable the positioning is. Some GPS manufacturers do provide a VDOP value for each record, but many only provide a more generic DOP value.

When σ_UERE_ and VDOP are known, the error-generating process can then be approximated by a Gaussian process with time-varying standard deviation σ_z_(t) = VDOP(t) · σ_UERE_ (Eq. 6.45 in Sanz Subirana et al., 2013). Therefore, the DOP is not a direct index of precision. The spread of the error distribution increases with the DOP, but the error on any given record is stochastic. The DOP is therefore not intended to be used as a data filter (e.g., discard any data with DOP above a given threshold), but instead it should be used to model the error-generating process.

### 2. Error in *z* when *z* is given by an altimeter

If recorded using an altimeter, *z* is computed from the barometric pressure, using the formula *z* = *c · T · log*(*P_REF_/P*) (Monaldo et al. 1986, Crocker and Jackson 2018). *c* is a calibration constant that mostly depends on the composition of the air (e.g., percentage of vapour) and on the gravitational field. *T* is the air temperature in Kelvin, *P* is the air pressure, and *P_REF_* is the air pressure at an elevation of reference (both pressures in mbar or in Pascal). However, this formula only holds when the atmosphere is at equilibrium. Changes in temperature, pressure, and air composition, i.e., the weather, alter the link between *z* and *P*. These influences are difficult to control fully because one would need to measure the weather variables both where the bird is, and at the reference elevation immediately below the bird. In other words, altimeters can be more accurate than GPS to monitor flight height, but only over short periods of time when the weather can be considered constant and the altimeter is calibrated for that weather. One should ideally regularly re-calibrate the altimeters using direct observations of flight height and accurate measures of *P_REF_* and *T*. Unfortunately, field calibrations are rarely feasible in practice (but see Shepard et al., 2016; Borkenhagen et al., 2018). The consequence is that altimeters are often miscalibrated. The degree of miscalibration depends mostly on the weather. This generates temporally autocorrelation in the error time series. Over a restricted time period, the error patterns are thus more akin to a bias (a systematic over- or under-estimation of flight height) than to an error in the statistical sense of a zero-mean, identically and independently distributed random process. Importantly, altimeter data still allow one to compute the derivative of flight height, i.e., climb rate, because the amount of bias can be considered constant over short periods of time. In Part 2.1, we will directly compare the errors from GPS and altimeters using controlled field experiments.

### 3. GPS horizontal error

(x,y) is also affected by a user equivalent range error and a dilution of precision (Fig. 1). The horizontal error in (*x, y*) can thus also be described as a Gaussian process with time-varying standard deviation: 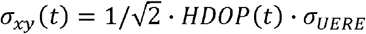. Note that we use here a horizontal dilution of precision factor, HDOP. An often-overlooked consequence of errors in the horizontal position is that they introduce flaws in the link to spatially-explicit environmental covariates (Hays et al. 2001, Bradshaw et al. 2007). In particular, the ground elevation *z_DEM_* is extracted from a location (*x, y*) that is slightly different from the true location (Katzner et al. 2012). If the terrain is very rough, then the ground elevation at the recorded location (*x, y*) may be significantly different from the ground elevation below the actual location of the bird. In Part 2.2 we will use simulations to quantify the influence of horizontal errors.

### 4. Interpolation error in *z_DEM_*

*z_DEM_* is interpolated from discrete ground elevation measurements (Gorokhovich and Voustianiouk 2006, Januchowski et al. 2010). The ground elevation is measured at a few select locations, but it is interpolated between them. The result of the interpolation is then rasterized at a set resolution, and the result is the DEM. This process can be quite imprecise (Gorokhovich and Voustianiouk 2006, Januchowski et al. 2010). At a cliff, for example, the ground elevation may drop by several hundred meters within a single pixel of the DEM.

### 5. Errors in DEM base data

The original measurements from which DEMs are interpolated are not necessarily error free either. These errors are assumed small relative to the other sources, however, there is, to our knowledge, not much information available about the base datasets from which DEM are interpolated and their precision.

## Part 2: Field trials, simulations, and reanalysis of raptor data

### Material and Methods

#### Controlled field trials

To quantify the magnitude of the vertical error in altimeters and GPS devices, we conducted three controlled trial experiments.

First, we attached an “Ornitrack 25” GPS-altimeter unit (Ornitela) to a drone. We then flew the drone above the rooftop of the Max-Planck institute in Radolfzell, Germany at heights ranging from 0 (drone landed on the rooftop) to 90m. We conducted 6 flight sessions over two days, each lasting between 15 and 140min, collecting one record every ten minutes for a total of 30 records. We also monitored the air pressure and temperature on the rooftop, which we used to recalibrate the altimeter post-hoc. Lastly, the drone carried a separate, on-board, altimeter.

In a second, separate experiment, we attached two “Gipsy 5” GPS units (Technosmart) to an ultra-light aircraft, with a vertical distance of 1.8m between the two units. We then flew the aircraft near Radolfzell while the two units simultaneously tracked its flight height, collecting one record per second for a total of 11.5 hours over 5 days.

Third, we compared the vertical positions recorded by 4 different units from 3 different manufacturers: Technosmart (AxyTrek and Gipsy 5), Microwave (GPS-GSM 20-70), and Ornitela (GPS-GSM Ornitrack 85). We (RG and OD) carried these units to 21 known geodesic points, of which the altitude was precisely documented by the French National Geographic Institute. The units recorded their position once every minute for a total of 894, 934, 560, and 563 data points, keeping only the unit * location combinations that yielded more than 25 fixes. We computed the bias and root mean squared error of the vertical measurement by comparing these data to the actual, known altitudes of the geodesic points. Importantly, the manufacturers do not use the same reference to compute the altitude: Microwave uses the geoid (WGS 84 EGM-96 norm), whereas the others use the mean sea level (assumed to correspond to the local reference, meaning the NGF-IGN 1969 norm, but sea below). We expressed all altitudes in the same norm before computing biases and errors, and accounted for sampling effort (number of fixes) and location when comparing the performance of different units.

#### Simulations of flight tracks

We simulated flight tracks that followed Ornstein-Uhlenbeck processes (Dunn and Gipson 1977). This is a class of continuous-time stochastic models, which is not specific to vertical movement or even to movement (Dunn and Gipson 1977). In the case of vertical movement, the parameters of the Ornstein-Uhlenbeck processes control the mean flight height, the variance in flight height, and the temporal autocorrelation in the flight height time series. We transformed the raw Ornstein-Uhlenbeck simulations using an atanh link as described by Péron et al. (2017) to enforce positive flight height. The time unit was arbitrary. An attractive feature of simulations in the context of this study is that we know both the *true* flight height and the *recorded* flight height, which is the true flight height plus an independent and identically distributed zero-mean Gaussian error.

#### Simulations of synthetic landscapes

The objective was to quantify the influence of horizontal errors. We generated synthetic landscapes of varying complexity and roughness (Fig. A2). We then transposed the flight track of a lesser kestrel *Falco naumanni* over these synthetic landscapes. The individual originally flew over extremely flat terrain (the Crau steppe in France). The data (Pilard and OD, unpublished) were collected every 3 minutes using a Gipsy 5 GPS unit from Technosmart, and processed through the state-space model of Péron et al. (2017) to account for real sampling errors before use. We then added simulated random telemetry noise of controlled standard deviation.

#### Raptor case studies

We reanalysed the data from Péron et al. (2017), where the field procedure, data selection, and data analysis procedures are described in full. Briefly, we studied three species of large soaring raptors: Andean condors *Vultur gryphus* (five juveniles, 1,692 individual.days of monitoring, 15 minute interval), Griffon vultures *Gyps fulvus* (eight adults, 2,697 individual.days, 1-5 minute interval), and Golden eagles *Aquila chrysaetos* (six adults, 3,103 individual.days, 6-10 minute interval). After applying the analytical procedure, for each data point, we could compare the *corrected* position, an estimate of the *true* position, to the *recorded* position, which was affected by the sources of errors we listed in Part 1.

We selected the period between 11:00 and 15:00, which concentrates condor activity and therefore flight time, and discarded other records. For the vultures, we selected the period between 09:00 and 16:00. For the eagles, we selected the period between 08:00 and 17:00 and, because a lot of time is spent motionless in this species even during their core activity period, we further removed all the records that were less than 15 meters from the previous record. We acknowledge the arbitrary nature of this data selection and emphasize that it is not necessary or even recommended to apply such filters before analysis. We however stress that in the context of the present study, the case studies perform an illustrative function, meaning that we use them to highlight the effect of improper error-handling, at least during the particular time periods that we selected for analysis because we consider them relevant for biological inference, and that the same analytical procedures can indiscriminately be applied to other time frames.

#### Collision risk

In several instances, we will illustrate the potential effect of improper data-handling on management recommendations by estimating the risk of collision with wind turbines as the proportion of records between 60 and 180m above ground (assuming no behavioural adjustment in the presence of wind turbines). Collision risk estimated from GPS tracks is increasingly used to make recommendations about the choice of locations for new turbines, or to schedule the operation of existing ones. We expected that the estimated collision risk would depend on flight parameters (mean flight height, variance in flight height), on the magnitude of errors, and on error-handling. For example, a large variance in flight height might lead to a high collision risk even if the mean flight height is beyond the collision zone. Improperly handled errors may lead to positions being erroneously recorded in the collision zone when the birds actually flew outside of it, and *vice versa*. The same type of thinking could be applied to other types of collision risk, e.g., antennas, utility lines, buildings with bay windows, except that the collision zone would be at a different height.

### Part 2.1: The magnitude of vertical errors in GPS and altimeters

During the first controlled field trial (with the drone), DOP values between 1.2 and 1.6 indicated that the configuration of the satellite network was reliable throughout. Nevertheless, 6.7% of the GPS flight height records were below the rooftop height, i.e., obviously erroneous. For the altimeter, with default settings, 10% of the records were below the rooftop height. The default settings of the altimeter therefore did not correspond to the atmospheric conditions during the experiment. The standard deviation of the difference between the recalibrated altimetry and the GPS data was 22m, between the recalibrated altimetry and default-setting altimetry it was 14m, and between the recalibrated altimetry and the on-board drone altimeter it was 19m. This means that, with default settings, the altimeters had approximately the same precision as the GPS.

During the second controlled field trial (with two GPS units attached to the same aircraft), in 35% of cases, the lower unit was erroneously recorded above the higher unit. The standard deviation of the difference between the height recorded by the two units was 7.1m. The highest of the two units recorded 3% of negative flight heights. The lowest unit recorded 13% of negative flight heights.

During the third controlled field trial (with GPS units carried to a geodesic point of precisely known altitude), the mean absolute bias of the vertical measurement was 27m on average across units and locations. The root mean squared error ranged from 14m to 42m depending on the unit, with a small effect of location. However, the within-session standard deviation ranged only to 28m, suggesting that a bias in the sea level reference point (probably incorrectly assumed to follow the French norm) inflated the RMSE. The average bias ranged between −17m and +12m depending on the unit, after correcting for significant location effect, but without effect of altitude. This means that different brands of GPS unit yield different rate of error in their altitude measurements, which can impair the comparison of datasets collected by different units. Further investigation or communication with manufacturers should decipher whether this stems from different fix acquisition procedures (e.g., satellite detection) or different post-processing algorithms, and should also make clear which sea level reference point different manufacturers are using.

These controlled field trials, along with other similar reports (Bouten et al., 2013; Ross-Smith et al., 2016), highlight that even in benign conditions, GPS and altimeter data are sufficiently error-prone to tamper with ecological inference in many cases (range of the standard deviation of the error: 4–50m). The issue is only suspected to be more acute in operational conditions when the DOP is larger, the terrain rougher, the weather more variable, and there are more obstacles to signal diffusion than in controlled field trials. Furthermore, the rate of error depended on the brand of the unit and on the location, which can be of importance when comparing across studies.

### Part 2.2: Horizontal errors can cause vertical errors

In the synthetic landscape simulations, the frequency of negative flight height records increased with the standard deviation of both the horizontal and vertical telemetry error (Fig. A2a), and with the landscape roughness and complexity (Fig. A2b). However, the various sources of errors acted in a multiplicative way, so that even when the telemetry noise was small (SD of 1m), the error in *h* could be large (SD of 20m; Fig. A2c; darkest grey curve). Perhaps unexpectedly, when the horizontal error was large, the error in the height above ground tended to be independent of the vertical error in the GPS (on average across all simulations; Fig. A2c; lightest grey curve). This means that the effect of the horizontal error in the GPS can supersede the effect of the vertical error, if the terrain is rough. Even in the absence of any vertical error, the horizontal error was indeed routinely sufficient to cause 10-20% of the data points to be below ground (Fig. A2a).

### Part. 2.3: Errors inflate the recorded variance in flight height

In the simulations of flight tracks, errors in *h* inflated the variance in the distribution of recorded flight heights, i.e., the variance in the true flight height was consistently lower than the variance in the recorded flight height (Fig. 2). In the raptor case studies, we obtained the same result, with the caveat that we did not access to the true flight height, but we could instead use the corrected flight heights (Fig. 2).

**Fig. 2:**
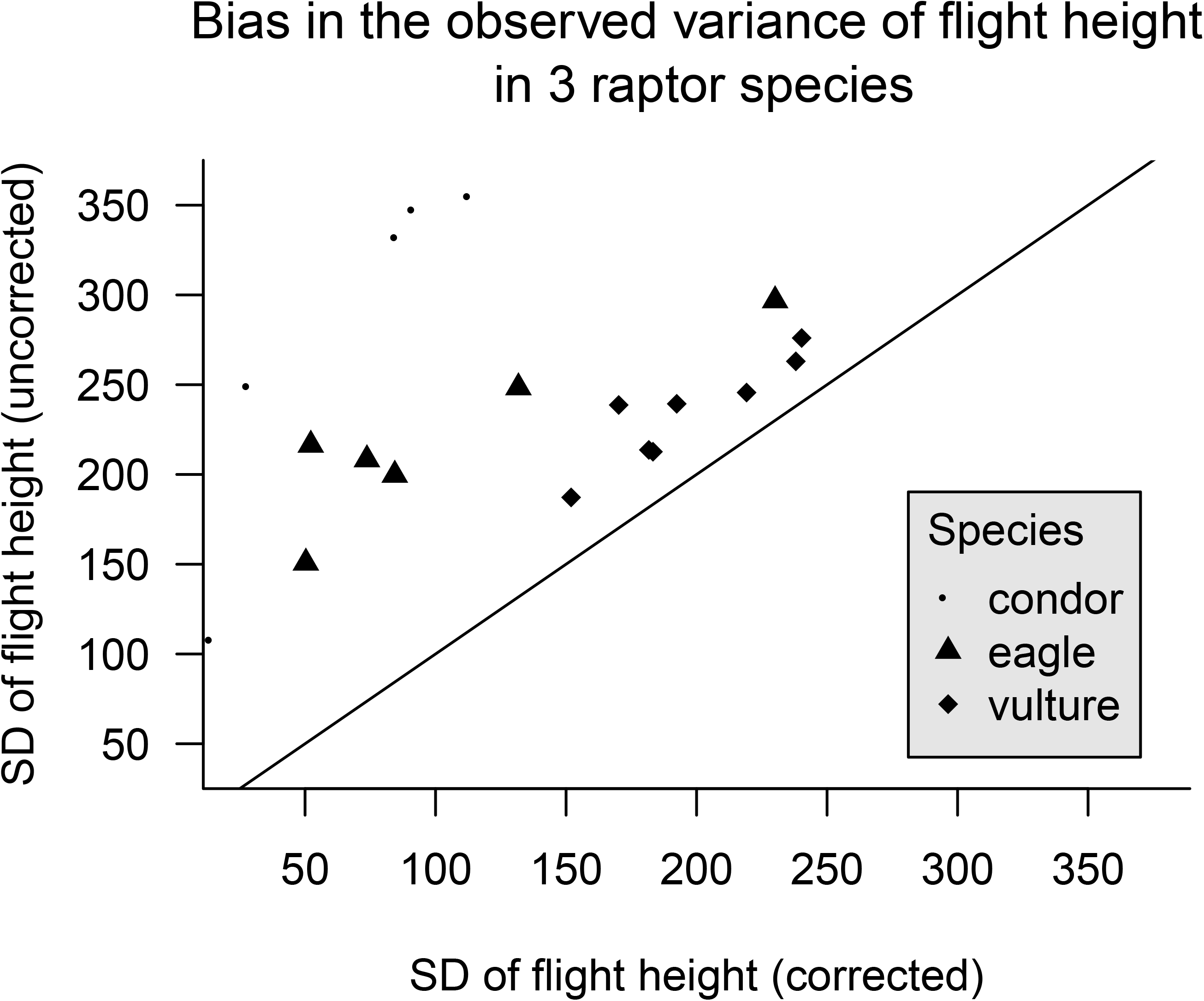
Comparison between the standard deviation of the *recorded* flight height (y-axis) and of the *corrected* flight height (x-axis), assumed to represent the *true* flight height, in three species of large soaring raptors. Each point stands for one bird over its entire monitoring period. The state-space model that we used to correct the flight heights, and in particular its robustness to variation in sampling resolution across populations, is explained in Péron et al. (2017). The diagonal line shows where the points should be if the recorded flight heights were error-free.

Indeed, if the movement and error processes are independent, the total variance in flight height is theoretically exactly the sum of the movement and sampling variances (e.g, Auger-Méthé et al., 2016; see also Gould & Nichols, 1998 and references therein). If the movement and error processes are not independent, the total variance is still larger than the movement variance. Yet, what we need for biological inference is the movement variance. In a naïve analysis of the raptor case studies that would confound telemetry errors with rapid movements, the birds would therefore have appeared more vertically mobile and with a more spread-out distribution in the aerosphere than they actually were. This type of issue is potentially quite widespread in movement ecology, e.g., in behavioural assignment exercises that use movement variances (daily displacements, turning angles, etc.) to determine the behavioural state of animals.

### Part 2.4: Negative flight height records provide useful information

In this section we focus on negative records, i.e., unrealistically low records, but the same logic can be applied to unrealistically high records. Negative flight height records are more likely to occur when animals are near the ground, either perched or flying. If we remove the negative records (Poessel et al. 2018), perching and low flight are under-sampled in the final dataset (Roeleke et al. 2018). To illustrate this point, we used a flight track from a migrating juvenile osprey (*Pandion haliaetus*) as it crossed the sea between the Italian mainland and Corsica (Duriez et al. 2018). During a portion of that sea crossing, its Ornitela GPS unit recorded flight heights that oscillated between −2m and −7m below the sea level (Fig. A3, inset). The amplitude of the oscillation suggested that the bird followed the swell of the waves. The complete sequence (Fig. A3) depicts a progressive loss of altitude as the bird glided towards firm ground, and a period of active flapping flight (as per the accelerometery record) very low above the waves once the bird had lost all of its accumulated potential energy before reaching firm ground. These negative flight height records documented a critical time period. First, the risk of having to make a sea landing were clearly much greater in the few minutes when the osprey was flying low over the waves, compared to the rest of the sea crossing when the bird was often soaring high (Duriez et al. 2018). In addition, when flying low, the bird had no other choice than to flap and therefore expend energy; whereas when higher above the sea, the bird had the option to soar and therefore spare energy. It is critical that negative flight height records like these are maintained, even if, instead of a fully interpretable high-resolution sequence like in this example, there are just a few isolated negative flight height records in a low-resolution dataset.

In addition, if we only kept the records with positive flight height, we would obtain a biased sample of the distribution of flight height. Both in simulations and in the raptor case studies, discarding negative flight height records led to the overestimation of the mean flight height in the remaining dataset, the underestimation of the variance in flight height, the introduction of a right skew in the distribution of flight height, and the overestimation of the collision risk (Fig. 3). The latter result was because negative records mostly occurred when the bird was flying below the collision zone, and thus removing negative records led to under-sample safe periods of time. Note that this particular result pertains to the wind turbine application case only; in other types of collision risk, e.g., buildings and utility lines, the collision zone starts closer to the ground.

**Fig. 3:**
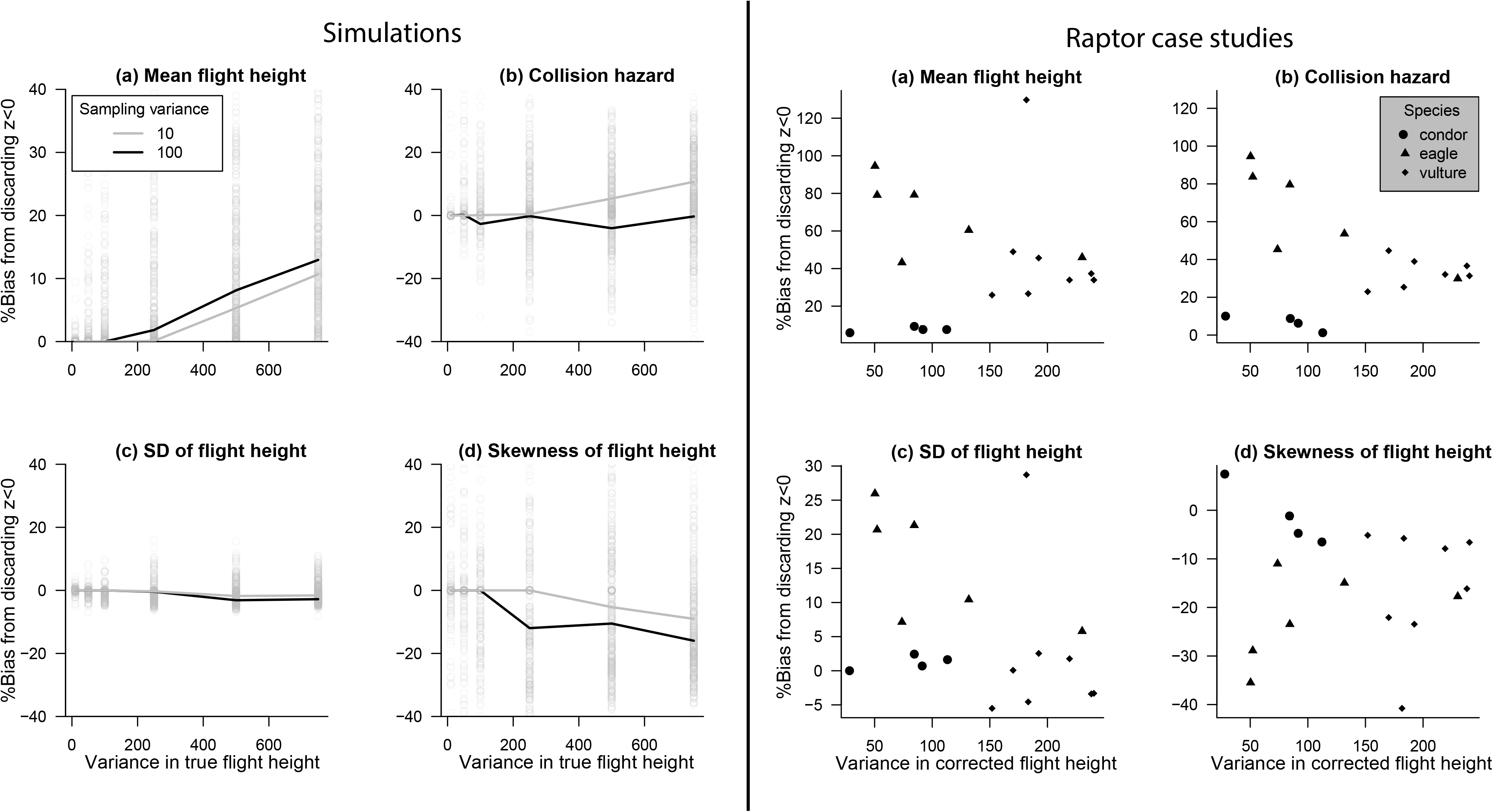
Removing the negative *recorded* flight heights introduces biases in the distribution of the remaining flight heights. Left group of panels: in simulations, where the *true* flight height is known. Right group of panels: in the raptor case studies, where the *corrected* flight height is assumed to represent the true flight height. In all panels, the x-axis features the variance in the true (or corrected) flight height. The y-axis features the percentage bias in (a) mean flight height; (b) collision risk (proportion of time spent between 60 and 180m above ground); (c) variance in flight height; and (d) skewness of the distribution of flight height. A percentage bias of +10% means that the focal quantity is 10% larger after we remove the negative records.

The simulations nicely complemented the raptor case studies by 1) eliminating any debate about whether the corrected flight heights in the raptor case studies were trustworthy or not (in the simulations, the true flight heights are exactly known) and 2) increasing the range of flight behaviours, since the raptors tended to exhibit lower percentage of time near the ground (in part because we purposely tried to exclude time spent perched) and different distributions of the sampling error. The amount of bias appeared highly dependent on the underlying flight behaviour and error distribution, and therefore not easy to predict and account for without appropriate error-handling methodology.

Additionally, there are many other major consequences of discarding negative flight heights. One is the disruption of the expected balance of positive and negative errors in the remaining data. Negative flight height records only arise when the error is negative, and so removing them introduces a bias towards positive errors, thereby disrupting the shape of the distribution of errors in the remaining data. Yet, we need the full range of errors to fit the models in Part 3. Another, unrelated consequence is the disruption of the sampling schedule of the remaining data. Many movement analyses are critically sensitive to the sampling schedule, and therefore their outcome will not be the same after removing the negative records. Lastly, and perhaps most importantly, negative flight height records can help fit the models that separate the error and movement processes, because they are unambiguously erroneous and can be informed as such in the model-fitting procedure (cf. Part 3). Some authors have applied less stringent filters, such as removing only the most negative flight height records and removing an equal amount of extremely positive flight height records. While the effect on the remaining distribution, and on the balance of negative and positive errors is supposedly weaker than if removing all of the negative records, we warn that the remaining records are still affected by the same error process that generated the records that were deemed too erroneous to keep, thus the issues in Part 2.3 still need to be addressed. In addition, these extremely erroneous records are potentially the most informative regarding the shape of the error distribution (cf. Part 3).

### Part 2.5: The mean flight height is not sufficient to describe the distribution of flight heights

Flight height datasets are often reduced to a single summary metric, the mean flight height and its variation with environmental and individual covariates (Walter et al. 2012, Cleasby et al. 2015, Poessel et al. 2018, Tikkanen et al. 2018, Balotari-Chiebao et al. 2018). This decision is mostly based on the ease of implementing spreadsheets, linear models, moving averages, or spline models. In this section we instead call for approaches that describe the full distribution of flight heights in the aerosphere, not only the mean flight height. To justify this call, we again focus on collision risk estimation. Indeed, if the variance in flight height is large enough, a proportion of time may be spent in the collision zone even if the mean flight height is outside the collision zone. In simulations, the proportion of time spent in the collision zone indeed depended on both the mean and the variance in flight height (Fig. 4a-b). In the raptor datasets, the estimated probability of flying in the collision zone did not decrease much for the individuals whose mean flight height was estimated above the collision zone (Fig. 4c). Similarly, the individuals that had an estimated mean flight height well below the collision zone were predicted to spend about 20% of their time in the collision zone (Fig. 4c). We strongly recommend that collision risk forecasts should not be based on the fixed effects of linear models, but instead on the full distribution of flight heights – a recommendation that will likely hold for all studies into vertical airspace use.

**Fig. 4.**
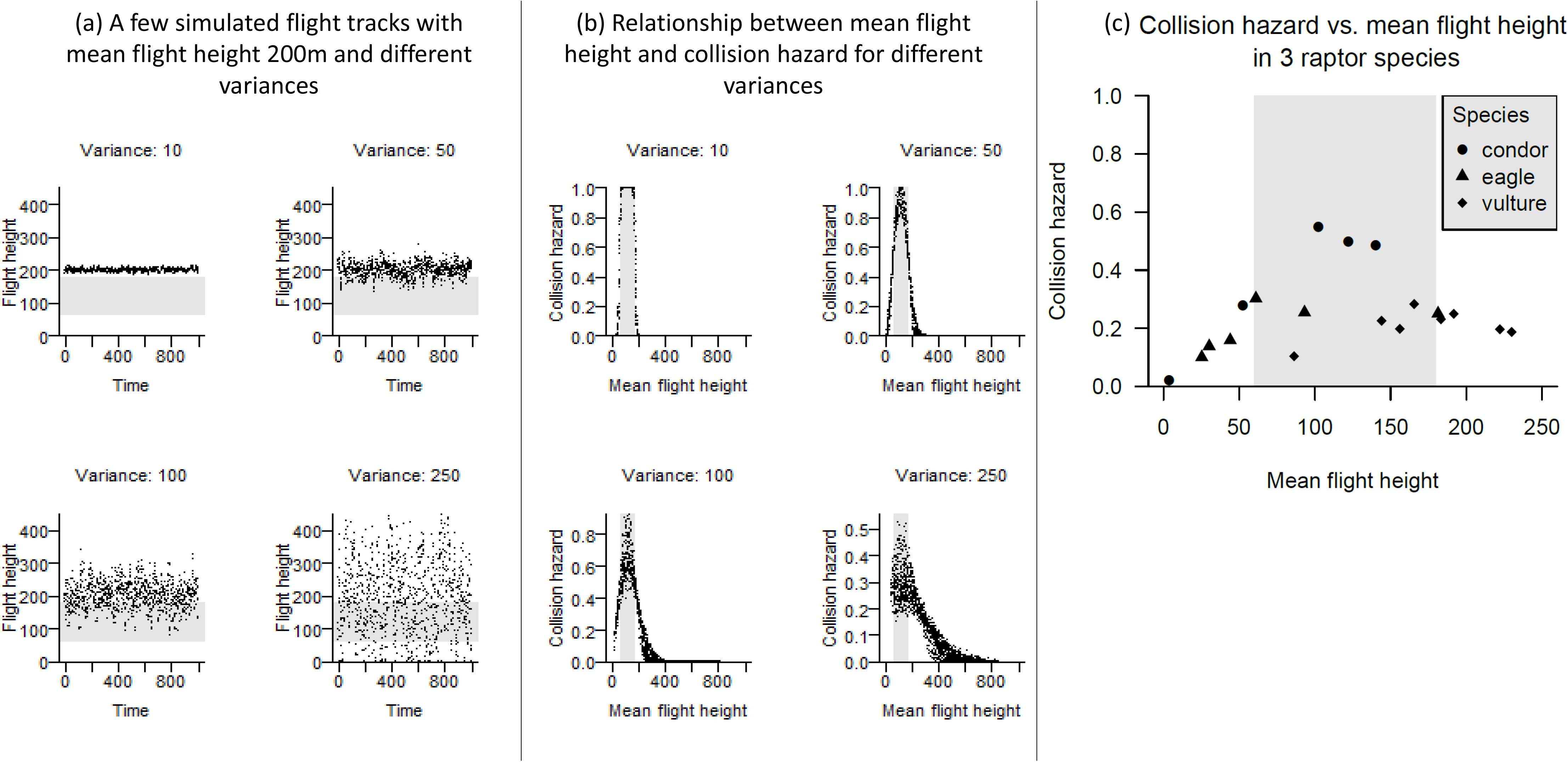
The variance in flight height influences the percentage of time spent in the collision zone of a wind farm (grey area, between 60 and 180 m). (a) Four simulated tracks (where the *true* flight height is known) with the same mean flight height (200m) but different variances (10, 50, 100, and 250m^2^). (b) More extensive simulations. Each point corresponds to one simulated track with a different mean flight height. (c) Same as (b) but using real datasets collected from three raptor species, where the *corrected* flight height is assumed to represent the true flight height. Each symbol stands for an individual over its entire monitoring period.

## Part 3: Statistical solutions

The state-space model framework (de Valpine & Hastings, 2002; Fig. 5) has a structure that is naturally aligned with the challenges of sampling errors in vertical space-use data. A state-space model is a stochastic model describing the changes over time in a state variable (here, the true flight height), when that variable is imperfectly observed (here, the recorded flight height). There is a “state process”, separated from an “observation process” (Fig. 5). State-space models are routinely used to correct for positioning errors in satellite-tracking data (chap. 6 in Sanz Subirana et al., 2013), including in wildlife studies (Patterson et al. 2008, Johnson et al. 2008, Albertsen et al. 2015, Brost et al. 2015, Buderman et al. 2015, Fleming et al. 2017). Importantly, these applications are not to be confused with another application of state-space models to movement data, when the focal state variable is a “behavioural state” whose Markovian transitions drive changes in movement rates (Gurarie et al. 2016, Pirotta et al. 2018, Murgatroyd et al. 2018). Instead, when the objective is to correct for positioning errors, the state variable is the position itself.

**Fig. 5:**
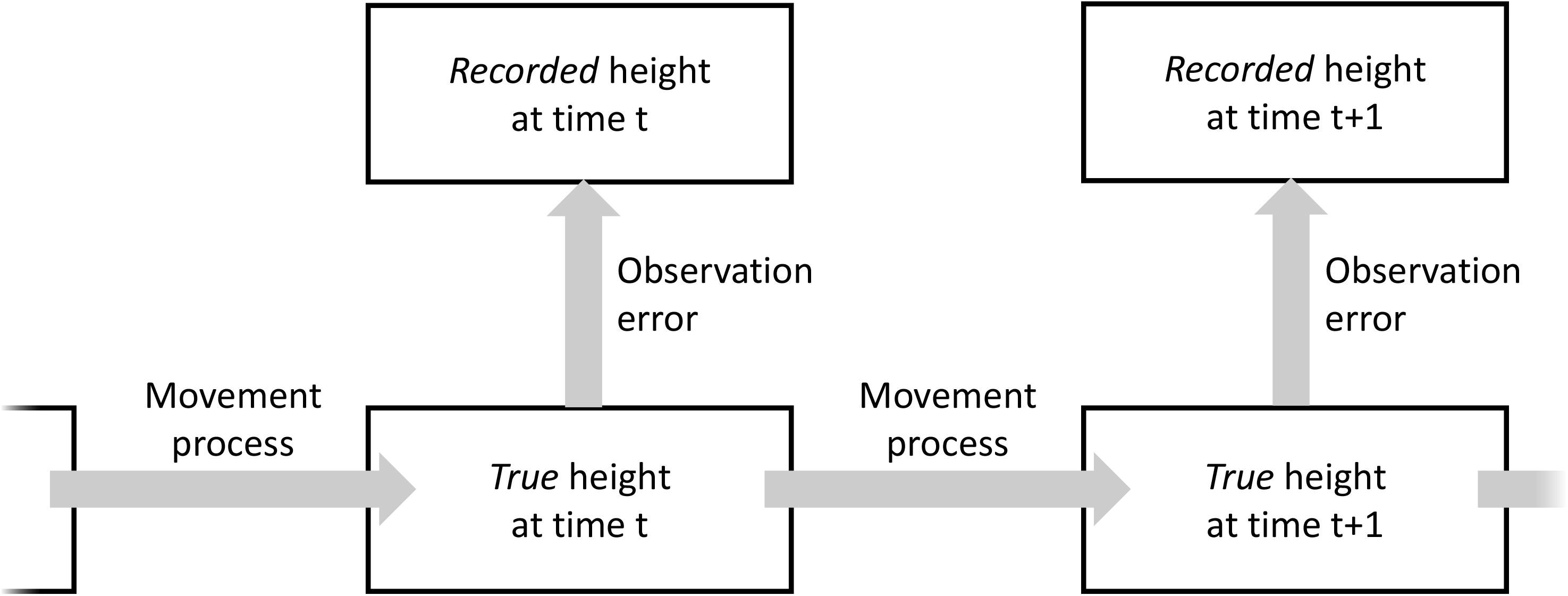
Schematic overview of the principles of a state-space model as applied to the correction of sampling errors in flight height data. The movement (or state) process accounts for the distribution of *true* flight heights. The observation process introduces sampling errors of various origins (Part 1) and yields the *recorded* flight heights. It also accounts for the sampling schedule. By fitting this model to recorded flight height time series, we can retrospectively compute the *corrected* flight height, an estimate of the true flight height.

In studies of flight height, the movement model can be set up such that the state variable always stays above zero. Then, if the recorded flight height is −7m, the model “knows” that the error was at least 7m (Ross-Smith et al., 2016; cf. Part 2.4). Actually, the presence of unambiguously erroneous records makes flight height studies better-suited to apply state-space models than many studies into horizontal space use by animals. Indeed, even when in theory the model is estimable, sometimes only a subset of the parameters of a state-space model are separately estimable, a phenomenon called “weak identifiability” that occurs when the sampling variance largely exceeds the process variance. An example of weak identifiability is when the difference between two classes of individuals are larger than the differences within the classes (Garrett and Zeger 2000). In addition, there are large statistical correlations between variance parameters in a movement model (Fleming et al. 2017), making it extra difficult to accurately separate movements and errors in sparse datasets. In that context, unambiguously erroneous records, such as negative flight heights, represent an additional source of information (Brost et al. 2015). They can help separate the process and sampling variances (Péron et al. 2017) and solve issues of weak identifiability.

As a perspective, we stress that there are also ways to obtain unambiguously correct records. These records could in theory perform a role similar to that of unambiguously erroneous records. For example, sometimes the position of the animals can be confirmed, e.g., at a documented feeding site, a nest, or by an incidental ground-based sighting. Those records can then be matched to the GPS track, yielding an exact measure of the local error. Animal-borne devices may also include a transponder designed to signal passage near strategically-placed emitters (e.g., Rebke, Coulson, Becker, & Vaupel, 2010). This type of validation data is routinely used in other applications of the GPS technology (Sanz Subirana et al. 2013). Lastly, the state-space framework is naturally conducive to the joint analysis of multiple sources of error-prone data (e.g., Péron, Nicolai, & Koons, 2012). In flight height studies, it is therefore possible to jointly analyse GPS and altimeter data, or multiple GPS streams coming from the same animal. This double-data approach is expected to help with statistical covariance issues, but cannot be expected to fully resolve all identifiability issues (Besbeas & Morgan, 2017), which only error-free validation data can do.

We should eventually stress that several wildlife GPS manufacturers already use a state-space model as part of the onboard data pre-processing algorithm, i.e., the released data have already been corrected by a proprietary state-space algorithm which may furthermore rely on proprietary validation data (Ornitela staff, pers. comm.). From our experience, in wildlife applications, these pre-processing algorithms are only applied during “bursts” of high-frequency data acquisition, not when the users request a more traditional low-frequency data acquisition schedule. Importantly, the data may not be pre-processed *across* bursts. The error from the first location of a burst is then carried over the entire burst sequence. Flight height tracks affected by this issue would exhibit a staircase-shaped profile. Overall, this type of data pre-processing trades a lower error variance against a larger error autocorrelation. Additional state-space modelling of the released pre-processed data can deal with this type of error autocorrelation, but the models need to be custom-made, i.e., are not routinely implemented in software. Perhaps more worryingly, some commercially-available GPS units apparently simply truncate the recorded height at zero above sea level (pers. obs.). We call for a more open approach to these data manipulations, including making the raw, unprocessed GPS records available, in addition to any pre-processed data, and with a formal description of the pre-processing algorithm.

We also acknowledge that the fitting of state-space models to vertical space use data still requires relatively rare statistical skills. Nevertheless, there are already several free, open-source computing environments to fit state-space models to vertical (and horizontal) movement data, and thereby estimate the most likely movement track as a by-product of the estimated parameters, similarly to how the individual values would be computed in a generalized mixed model with individual random effects:

- The crawl (Johnson et al. 2008) and ctmm (Calabrese et al. 2016) packages for R. These compute the likelihood of the state-space model using a Kalman filter. This algorithm is fast but requires all the model processes to be Gaussian or approximately Gaussian (no truncation or constraint, no excess extreme values, no excess kurtosis or skew).
- The TMB package for R (Kristensen et al. 2014) approximates the likelihood of the state-space model using the automatic differentiation algorithm with Laplace approximation. That approach makes computing times shorter than the next option, while still allowing for flexible modelling such as non-Gaussian errors (Albertsen et al. 2015), custom link functions (Péron et al. 2017), or multiple data streams.
- The Monte Carlo Markov Chain Bayesian framework (Plummer 2003, Spiegelhalter et al. 2003, Csilléry et al. 2010) generates parameter distributions that iteratively converge towards the solution. This option is the most flexible in terms of nonlinearities and non-Gaussian features, such as truncated distributions (Brost et al. 2015), but the computing time can be prohibitive large for datasets.

## Conclusion

Improper error-handling methodologies yield a flawed picture of aerial niches. For example, discarding negative flight height records artificially truncates the observed distribution of flight heights (Fig. 3), and focusing on the mean flight height alone (for example when using linear models) does not fully describe the aerial niche (Fig. 4). While these observations are quite intuitive, bad practices remain common enough that it was important to stress these issues and illustrate them thoroughly. On the other hand, not addressing the occurrence of errors at all would artificially spread-out the observed distribution of flight heights (Fig. 2), leading for example to increased observed vertical overlap between species and individuals, which can modify the inference about community processes. Improper error handling procedures would also tamper with the quantification of behaviour and flight strategies, by increasing or decreasing the observed vertical velocity, and interfere with behavioural state assignments. Lastly, errors may covary with environmental covariates such as terrain roughness and wind speed, e.g., GPS positioning precision decreases with terrain roughness (D’Eon et al. 2002) and wind speed decreases near the ground (Sachs 2005). Thereby, selectively discarding records based on the number of available satellites or the dilution of precision would lead to imbalanced sampling of terrain roughness, and discarding negative flight height records (that predominantly occur near the ground) would lead to misrepresent the relationship to wind speed.

Regarding applied consequences, we focused on demonstrating how improper methods would imperfectly quantify the time spent by GPS-tracked raptors in the rotor-swept zone of wind turbines (Fig. 3b). There are many other human-wildlife conflicts for the use of the aerosphere, for example bird strikes near airports and disturbance of wildlife by drones and other recreational aircraft. Regarding bird strikes, GPS-based predictive models of bird flight height (e.g., Péron et al. 2017) might help plan ahead the operation of airports. The state-space class of model that we advocate is actually already used, in real time, to exploit bird activity data from radar monitors and generate a warning system for airport managers (Bruder 1997). Regarding recreational aircraft and drones, analysing bird-borne GPS tracks may help reveal the effect of the disturbance, which is expected to increase in frequency as drones in particular become more popular (Rebolo-Ifrán et al. 2019). The recommendations we made about the effect of errors on the estimation of aerial niche overlaps and the quantification of behaviours seem particularly relevant in this context.

In conclusion, the issue of properly handling errors in flight height data is key to any aeroecology study. We strongly advise against ad-hoc “data quality” filters, and against statistical tools that only document variation in the mean flight height instead of the full distribution of flight height. Our proposed statistical framework based on state-space models and the analysis of the full distribution of flight heights requires interdisciplinary work between experts in flight behaviour and experts in data analysis, and the emergence of interface specialists, but the insights and the applied decisions based on those insights are expected to be more reliable.

## List of abbreviation

*h*: flight height above ground;
*z*: flight altitude (relative to the same reference as the DEM, e.g., the ellipsoid);
DEM: digital elevation model;
UERE: user equivalent range error;
DOP: dilution of precision;
SD: standard deviation.

## Declarations

### Ethics approval and consent to participate

Permissions to trap and tag griffon vultures and lesser kestrels were given by the Centre de Recherche sur la Biologie des Populations d’Oiseaux, Museum National d’Histoire Naturelle, Paris to 0. Duriez (Personal Program PP961) and to P. Pilard (PP311). Permissions to trap and tag osprey were given by Instituto Nazionale per la Protezione e la Ricerca Ambientale (ISPRA) under the authorization of Law 157/1992 [Art. 4 (1) and Art. 7 (5)] and under authorization no. 2502 05.05.2016 and no. 4254 27.03.2018 issued by Regione Toscana. Permissions to trap and tag condors were given by the Dirección de Fauna de Río Negro (DFRN), the Administración de Parques Nacionales (APN) from Argentina and the owners and managers of local farms. All condor procedures were approved by the DFRN, RN132.730-DF, and APN, 14/2011.

### Consent for publication

Not applicable

### Availability of data and material

The data have been uploaded to MoveBank (www.movebank.org)

### Authors’ contributions

All authors conceived the project and revised the manuscript. GP performed the analyses and simulations and wrote the first draft of the manuscript. OD, RG, SAL, KS, and ES procured the data.

## Acknowledgements

A. Scharf and H. Wehner (MPI) flew the drone and assisted with data collection. M. Quetting flew the ultra-light aircraft and E. Lempidakis assisted with data collection. We warmly thank everyone involved in the collection and management of the raptor tracking data, and in particular C. Itty for the eagle data, and the organizations *Asters* and *Vautours des Baronnies* for the help with field experiments.

## Supplementary material

Appendix A: Supplementary figures A1, A2, A3.

