## Appendix S1 for "The challenges of estimating the distribution of flight heights from telemetry or altimetry data"

**Supplementary figures**

Fig. S1: Schematic review of the sources of error in flight height data. Notation as in the main text. Circled numbers refer to subheadings in Section 1. Inset 1: Simplified representation of the role of UERE and DOP in the error of GPS localisations, in a favourable (left) and unfavourable (right) configuration of a two-satellite network.

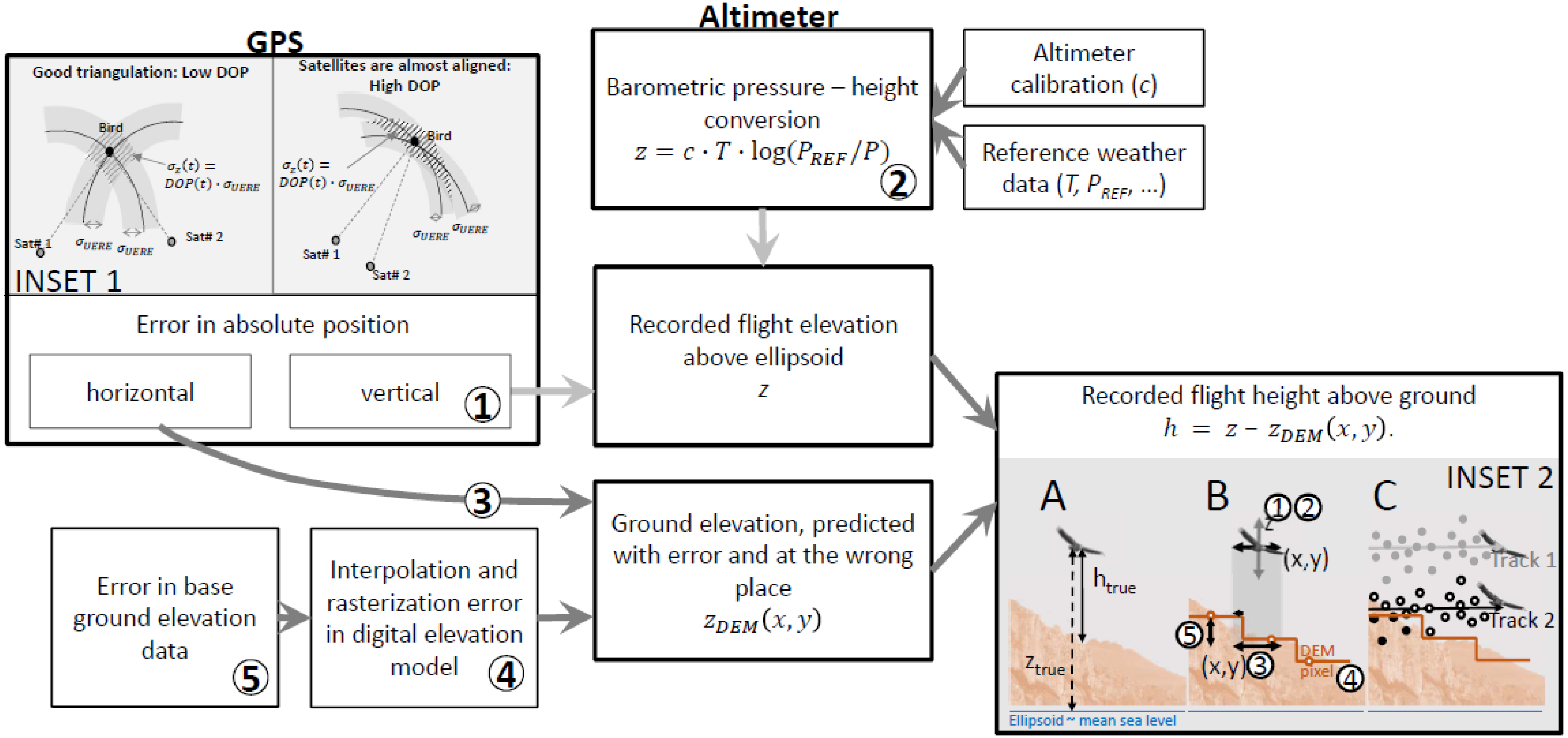

Fig. S2: The horizontal error of the GPS can create vertical errors in recorded flight height above ground. (a) 3D representation of the principles of the simulations. The true flight height is transposed over a simulated landscape, and thus initially there are no records below the ground elevation. We add telemetry noise both horizontally and vertically (standard deviation of the noise in this example: 10m). The vertical lines indicate the records that are now shifted below the ground elevation (17% in this example). (b-e) Analysis of 7000 similar simulations, changing the landscape structure and the amount of horizontal and vertical noise. (b) Percentage of negative flight height records as a function of the observation noise (vertical and horizontal). (c) Percentage of negative flight height records as a function of landscape structure (complexity and roughness, which control the "spikiness" of the landscape). (d) Standard error of the difference between recorded and true flight height as a function of the observation noise (vertical and horizontal). (e) Standard error of the difference between recorded and true flight height as a function of landscape structure (complexity and roughness).

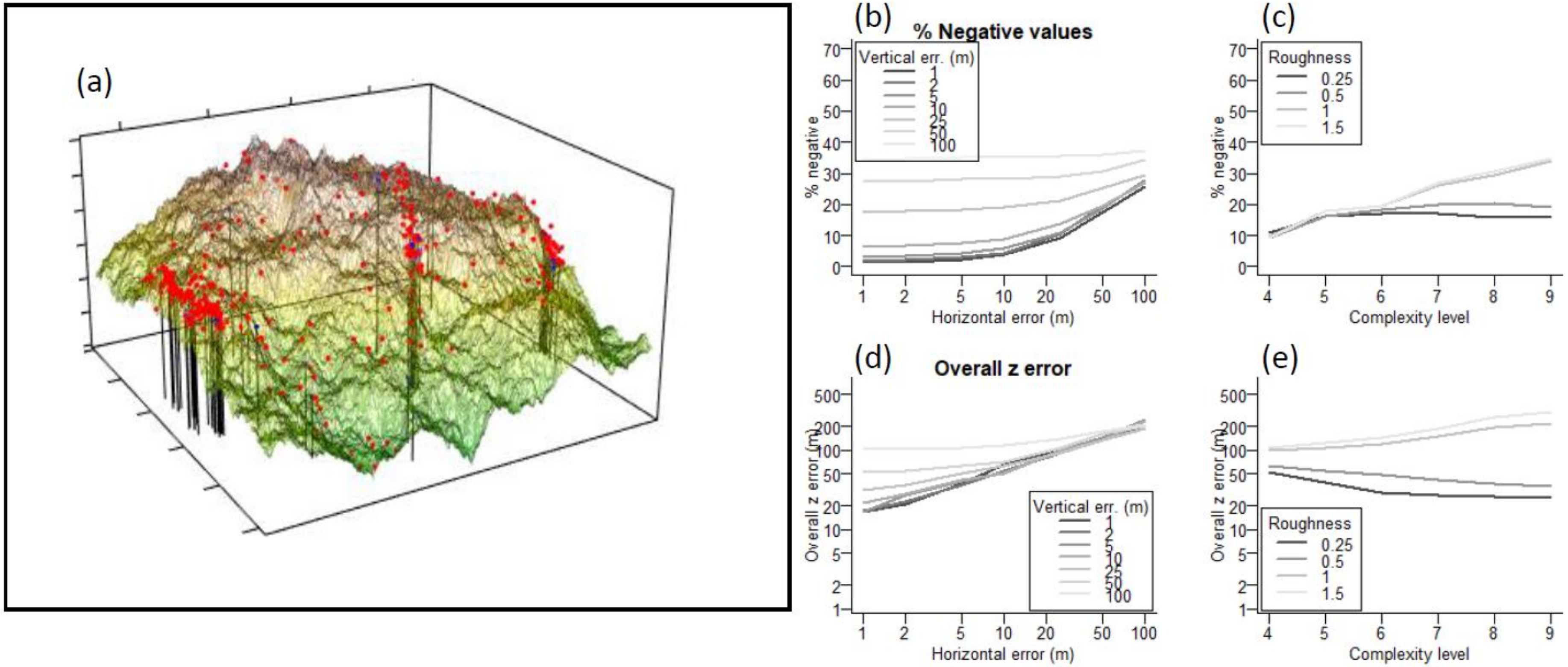

Fig. S3: 30 minutes in the life of a migrating osprey. Negative flight height records (red symbols, below the sea level) occur when the bird is flying low above the sea. Negative records document a biologically significant period of time when the bird had no potential energy left and was at risk of having to make a sea landing. The inset represents the raw data (grey symbols) and a moving average suggesting that the bird was closely following the swell of the waves. Imagery: Google Earth.

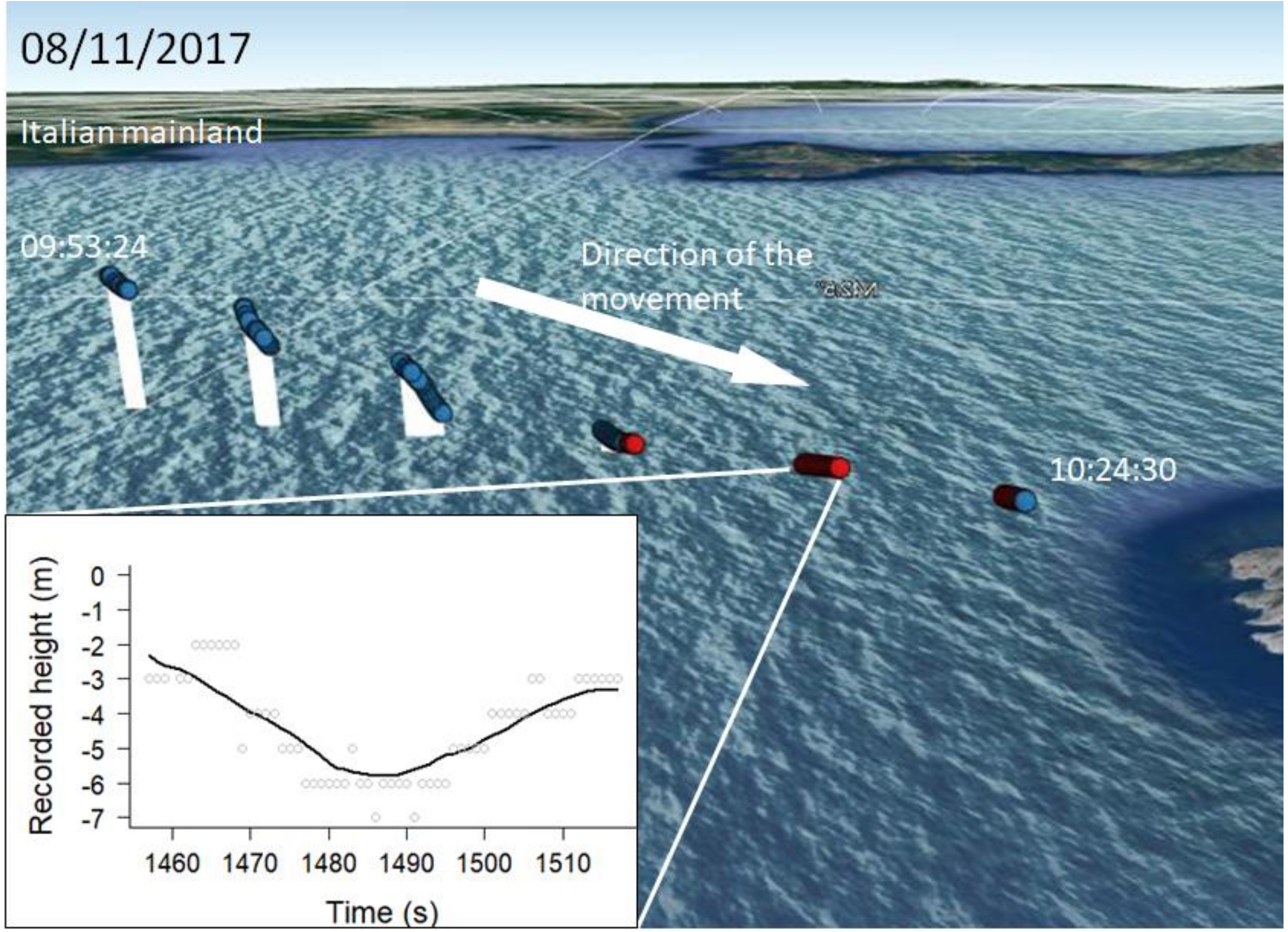
